# When time really is money: in situ quantification of the strobilurin resistance mutation G143A in the wheat pathogen Blumeria graminis f. sp. tritici

**DOI:** 10.1101/2020.08.20.258921

**Authors:** Kejal N Dodhia, Belinda A Cox, Richard P Oliver, Francisco J Lopez-Ruiz

**Affiliations:** Centre for Crop and Disease Management, Curtin University, 6102 Bentley, Australia; School of Molecular and Life Sciences, Curtin University, 6102 Bentley, Australia

**Keywords:** fungicide resistance, strobilurins, QoI fungicides, *Blumeria graminis* f. sp. *tritici* (wheat powdery mildew), allele specific qPCR, digital PCR, Cytb, in-field quantification, G143A

## Abstract

**Background:** There has been an inexorable increase in the incidence of fungicide resistance in plant pathogens in recent years. Control of diseases and the management of resistance would be greatly aided by rapid diagnostic methods. Quantitative allele specific PCR (ASqPCR) is an ideal technique for the analysis of fungicide resistance in the field as it can both detect and quantify the frequency of mutations associated with fungicide resistance. We have applied this technique to the fungal pathogen *Blumeria graminis* f. sp. *tritici* (*Bgt*), an obligate biotrophic fungus that causes wheat powdery mildew and is responsible for up to 25% yield loss annually. In Australia, strobilurin resistant *Bgt* was first discovered in samples from Tasmania and Victoria in 2016. Molecular analysis revealed a nucleotide transversion in the cytochrome *bc1 enzyme (cytb) complex*, resulting in a substitution of alanine for glycine at position 143 (G143A) in Cytb.

**Results:** We have developed an in-field ASqPCR assay that can quantify both the resistant (A143) and sensitive (G143) *cytb* alleles down to 1.67% in host and *Bgt* DNA mixtures within 90 min of sample collection. The *in situ* analysis of field samples collected during a survey in Tasmania revealed A143 frequencies ranging between 9-100%. We validated the analysis with a newly developed laboratory based digital PCR assay and found no significant differences between the two methods.

**Conclusion:** We have successfully developed an in-field quantification method, for a QoI resistant allele, by pairing an ASqPCR assay on a lightweight qPCR instrument with a quick DNA extraction method. The deployment of this type of methodologies in the field can contribute to the effective in-season management of fungicide resistance.

## Introduction

One of the major challenges that the agricultural industry faces is the control of fungal crop diseases during the emergence of fungicide resistance.^1–4^ The presence of fungicide resistant pathogen populations in a crop not only results in lower efficacy of chemical control methods but also lower yields and quality due to increased disease pressure.^5^ An optimal approach to combat this would be to determine what chemistries the populations are resistant to and use other, more efficacious chemicals instead.^2^

Many cases of fungicide resistance have been functionally linked to specific genotype changes in target genes. The detection of these genotypic changes is a valuable complement to the phenotypic screening of suspect pathogen isolates. Phenotypic screening requires the isolation of numerous pure cultures of the pathogen and testing for quantitative growth inhibition. Such assays are very demanding of space, resources and time, taking many days or weeks. The potential of screening using genotypic methods has been recognised for more than 25 years.^6^ A wide range of techniques have been used but all are limited by the need to isolate pure cultures of the pathogen and to carry out the analyses in a laboratory environment. As such the minimum time for such an assay is still several days. Therefore, the time-frame within which to spray a fungicide can be lost by the time the samples are sent for analysis to a laboratory and results received.

Other methodologies such as loop mediated isothermal amplification (LAMP) are rapid and can be used with infected plant samples.^7–9^ Based on this, LAMP has been used for the successful in-field and laboratory detection of different organisms including chemical resistance in fungi and weeds.^10–14^ However, the results from this test are not quantitative and prone to false positives.

An ideal solution to this will be the use of an assay, which can be performed rapidly, *in situ,* with high confidence results, to quantify the population of fungal pathogens that are resistant to a given fungicide. This would prevent the spraying of a fungicide to which a fungus is resistant, as that would both fail to control the disease and increases the selection of fungicide resistant populations.^15^ A rapid assay diagnosing the disease, its quantity and fungicide sensitivity status would allow farmers to choose the optimum fungicide regime for on-going protection of the crop.

Allele specific polymerase chain reaction (ASPCR)^16^ is a specific and sensitive assay that was developed for detecting point mutations using PCR followed by gel electrophoresis. This method has been adapted for use with both intercalating dyes and probe based assays to quantify fungi with genetic changes correlated with fungicide resistance.^17–19^ If combined with a crude DNA extraction and a ‘fast’ and robust polymerase that can tolerate field conditions, this assay could be conducted in the field with minimal laboratory equipment.

*Blumeria graminis* [DC] f. sp. *tritici* E.O. Speer (synonym *Erysiphe* graminis DC Em. Marchal) (*Bgt*) is an obligate biotrophic pathogen of wheat (*Triticum aestivum* L.) causing wheat powdery mildew (WPM).^20^ *Bgt* is often controlled with strobilurins or quinone outside inhibitors (QoI), which target the cytochrome *bc*_1_ enzyme (*cytb*) complex.^21,22^ The combination of its large population size and intensive control using fungicides has enabled the selection of genotypes with resistance to this group of chemicals. Strobilurin resistant genotypes of *Bgt* were first discovered in 1998 in Germany, just two years after their introduction for the control of various crop diseases, and have since been reported in wheat growing regions around the world.^21,23,24^ Sequencing of the *cytb* gene from *Bgt* showed that a single nucleotide polymorphism (SNP) of a guanidine to cytosine at position 428 (*g428c*), was the only difference between the sensitive and resistant isolates. ^21^ This mutation results in the amino acid change of glycine to alanine at position 143 (G143A) in the Cytb protein, (homologous to the archetypal *G143A* mutation in the Cytb of *Zymoseptoria tritici*).^21,25^

In Australia, over 10 Mha are sown with wheat annually, thus providing a large breeding ground for this pathogen.^26^ However, due to the range of climatic zones within the Australian wheat belt, mildew infections are most common in areas with medium to high rainfall as cool humid conditions are required for optimal disease development. The first report of strobilurin field resistance in Australian *Bgt* was from the states of Victoria and Tasmania in 2016 (F. Lopez-Ruiz, personal communication).

Here we report the development of a quantitative allele specific PCR (ASqPCR), performed on a portable thermal cycler, which can be powered by a battery, coupled with a simple 2-step DNA extraction protocol. The method successfully generated accurate genotype frequencies of the *B. graminis* f. sp. *tritici* Cytb G143A fungicide resistance mutation, on site, within 90 min of sample collection. To our knowledge, this is the first *in situ* ‘closed tube’ allele quantification in a plant pathogen.

## Materials and Methods

### Blumeria graminis tritici (Bgt) field sample PCR and cytb sequencing

Ten wheat and one barley powdery mildew samples were collected from the states of Western Australia, South Australia, Victoria and Tasmania in 2016 (Table 1). Infected tissue was placed in 15 ml polypropylene tubes filled with 2 ml of 50 mgL^−1^ benzimidazole water agar,^27^ sealed and placed in polystyrene containers with ice packs for express shipment to the laboratory for DNA extraction. For each sample, approximately half of the infected tissue was cut into small pieces and DNA was extracted using a BioSprint 15 instrument and BioSprint 15 DNA plant kit (Qiagen, Australia) according to the manufacturer’s protocol. The resulting mix of fungal and wheat DNA was eluted in 1 × TE buffer pH 8.0 and stored at −20 °C. The remaining sample was stored at −20 °C until further analysis.

**Table 1.**
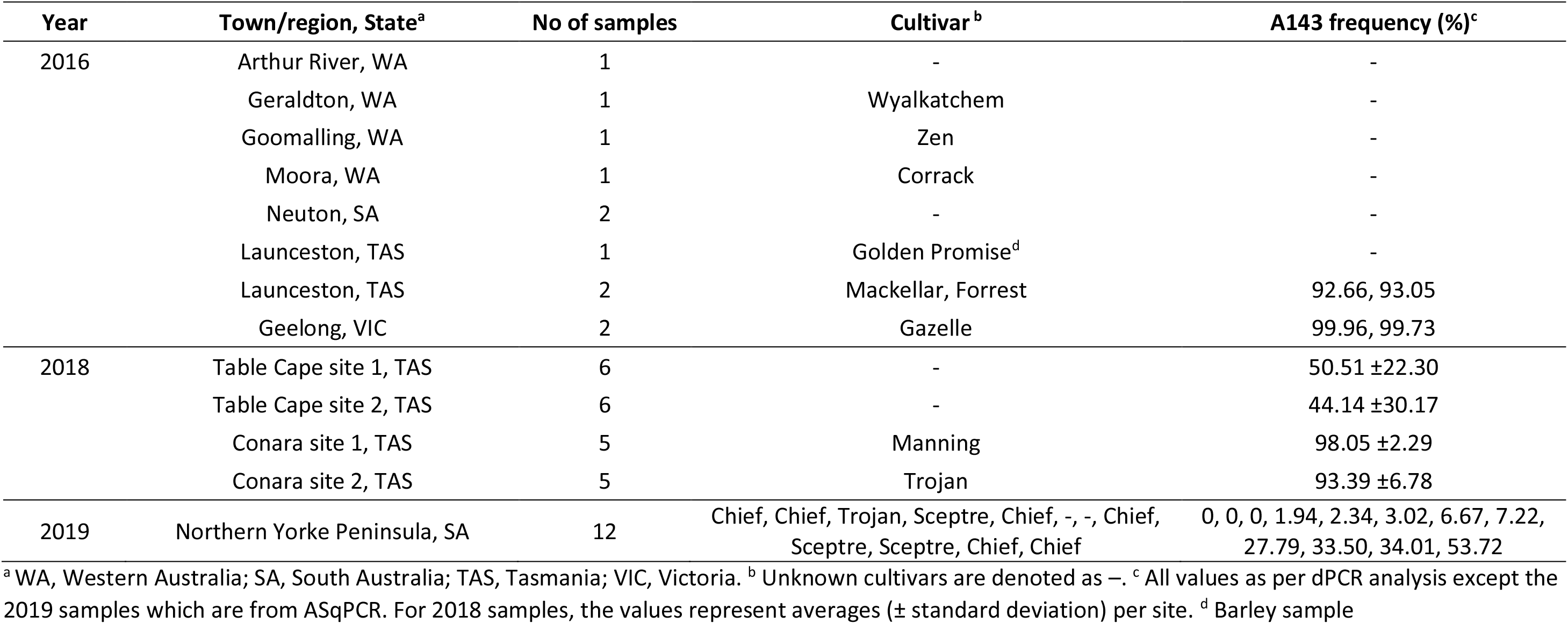
Sample details and *cytb* A143 allele frequency of wheat infected with *Blumeria graminis* f.sp. *tritici* collected in this study.

| Year | Town/region, State<sup>a</sup> | No of samples | Cultivar<sup>b</sup> | A143 frequency (%)<sup>c</sup> |
| --- | --- | --- | --- | --- |
| 2016 | Arthur River, WA | 1 | - | - |
|  | Geraldton, WA | 1 | Wyalkatchem | - |
|  | Goomalling, WA | 1 | Zen | - |
|  | Moora, WA | 1 | Corrack | - |
|  | Neuton, SA | 2 | - | - |
|  | Launceston, TAS | 1 | Golden Promise<sup>d</sup> | - |
|  | Launceston, TAS | 2 | Mackellar, Forrest | 92.66, 93.05 |
|  | Geelong, VIC | 2 | Gazelle | 99.96, 99.73 |
| 2018 | Table Cape site 1, TAS | 6 | - | 50.51 ±22.30 |
|  | Table Cape site 2, TAS | 6 | - | 44.14 ±30.17 |
|  | Conara site 1, TAS | 5 | Manning | 98.05 ±2.29 |
|  | Conara site 2, TAS | 5 | Trojan | 93.39 ±6.78 |
| 2019 | Northern Yorke Peninsula, SA | 12 | Chief, Chief, Trojan, Sceptre, Chief, -, -, Chief, Sceptre, Sceptre, Chief, Chief | 0, 0, 0, 1.94, 2.34, 3.02, 6.67, 7.22, 27.79, 33.50, 34.01, 53.72 |

<sup>a</sup> WA, Western Australia; SA, South Australia; TAS, Tasmania; VIC, Victoria. <sup>b</sup> Unknown cultivars are denoted as -. <sup>c</sup> All values as per dPCR analysis except the 2019 samples which are from ASqPCR. For 2018 samples, the values represent averages (± standard deviation) per site. <sup>d</sup> Barley sample

The *cytb* gene from *Bgt* was amplified and sequenced using primers WM-Cb-f and WM-Cb-R^28^ (Table 2). Each 100 μl PCR reaction contained 5 U MyTaq DNA polymerase (Bioline, Australia), 1 × MyTaq reaction buffer, 0.4 μM of each primer, DNA template and water. The thermal-cycling was conducted with initial denaturation for 5 min at 95 °C, followed by 34 cycles of 30 s at 95 °C, 1 min at 55 °C and 1 min at 72 °C, followed by final extension for 10 min at 72 °C. The PCR products were sequenced by Macrogen (Seoul, South Korea) and the results aligned on Geneious (Biomatters Ltd) with existing *Bgt cytb* gene reference sequences (GenBank accessions: AF343441.1, AF343442.1) to determine if mutation *g428c,* responsible for amino acid substitution G143A, was present.

**Table 2.**
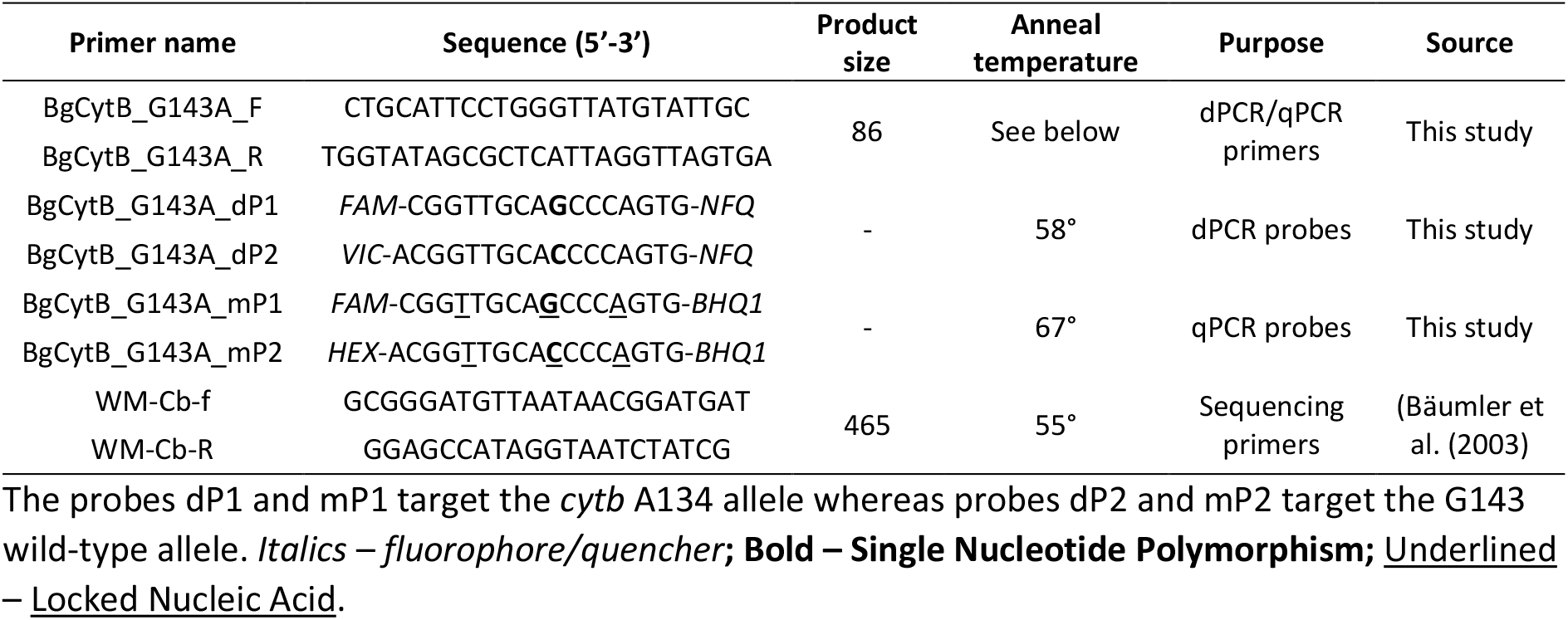
PCR, Digital PCR and qPCR primers and probes used in this study.

### Design of digital PCR and ASqPCR for the analysis of g428c

The digital PCR (dPCR) primers and TaqMan^®^ probes used to quantify the presence of mutation *g428c* in this study were designed as per Zulak *et al.* ^29^ with some modifications (Table 2). The sequence of *cytb* flanking 100 bp of the mutation was added to the Custom TaqMan^®^ Assay Design Tool, with the species/scale set at ‘Non-Human: small scale’. Resulting *Bgt cytb* wild type (G143 allele; strobilurin-sensitive) and mutant (A143 allele; strobilurin-resistant) primers and probe combinations were ordered as a pre-mix (Thermo Fisher Scientific, Australia). The probe targeting A143 allele was labelled with the fluorophore FAM on its 5’ end and the probe targeting the G143 allele labelled with VIC. The 3’ ends of both probes were labelled with a non-fluorescing quencher (NFQ).

Each dPCR reaction consisted of 8.5 μl QuantStudio™ 3D Digital PCR 2 ×Master Mix (Applied Biosystems, Australia), 0.3 μl primer probe pre-mix, made up to 17 μl with DNA template and water. 15 μl of this reaction was loaded onto a QuantStudio™ 3D Digital PCR Chip v2 (Applied Biosystems) and cycled on a Geneamp 9700 flat block thermal cycler (Applied Biosystems) under the following conditions: 10 min at 96 °C, then 40 cycles of 2 min at 58 °C and 30 s at 98 °C, followed by 2 min at 58 °C, and 10 min at 22 °C. Chips were read on a QuantStudio™ 3D Digital PCR Instrument (Applied Biosystems). The copy ratio of mutant to wild type allele in each sample was determined using QuantStudio™ 3D AnalysisSuite™ software (Applied Biosystems). The DNA from the sequenced samples was tested along with DNA from uninfected wheat leaf, wheat leaf infected with *Puccinia triticina* and *Parastagonospora nodorum* fungal DNA. In addition, a dilution series of the DNA from the Goomalling sample (G143 allele, table 1) and Geelong 1 sample (A143 allele, table 1) was tested with their respective target probes to determine the detection limits of the assay. Each test was conducted in triplicate.

dPCR primers and modified dPCR probes (Sigma Aldrich, Australia) were used to detect mutation *g428c* on a magnetic induction cycler (MIC) qPCR instrument (Biomolecular Systems, Australia). In order to increase probe *T*_*m*_ and specificity, three bases in each probe were modified to Locked Nucleic Acids (LNA) (Table 2). In addition, the fluorophore on the probe targeting the G143 allele was changed from VIC to HEX.

Three different qPCR mixes were tested using the Goomalling and Geelong 1 DNA (Table 1). Each 20 μl ASqPCR reaction consisted of 10 μl of 2x mastermix (iQ Multilplex Powermix (Bio-Rad, Australia), Sensifast Probe No-Rox mastermix (Bioline) or ImmoMix™ (Bioline)), 0.5 μl each of 10 μM forward and reverse primers, 0.3 μl each of 5 μM FAM and HEX probes, 5 μl DNA template and 3.4 μl water. The assay was conducted under field conditions using the following conditions: initial denaturation for 5 min at 95 °C, then 40 cycles of 10 s at 95 °C, and 30 s at 67 °C for annealing and extension. Each sample was replicated five times. The Sensifast Probe No-Rox mastermix (Bioline, Australia) was chosen for future testing.

A dilution series similar to that described for dPCR including single and mixture of known allele proportions was tested to determine the detection limits of the assay. A standard curve was also constructed using a serial dilution of the Goomalling and Geelong 1 DNA to calculate the amount of fungal DNA in the mixtures. The A143 allele frequency was calculated as percentage value using MIC software v2.8.0 (Biomolecular Systems). The species specificity of the assay was tested using DNA from plant material infected with closely related pathogens *Blumeria graminis* f. sp. *hordei* and *Erysiphe necator*, as well as fungal DNA from pathogens sharing the same host; *P. nodorum* and *Pyrenophora tritici-repentis*.

### Development of an in-field allele frequency quantification pipeline

To enable the testing to be conducted in a field setting, a quick DNA extraction protocol was developed as a part of this study. Approximately 100 mg of infected plant tissue that had been kept at −20 °C were crushed with a plastic micro pestle in 400 μl of the lysis buffer (0.5% w/v SDS, 1.5% w/v NaCl) until the solution appeared green and the plant tissue fragmented. This was then diluted 100-fold in 1x TE buffer pH 8.0, 5 μl of which was used as DNA template in ASqPCR reactions. DNA extractions were conducted under field conditions, and analysed using the cycling protocol described previously.

Twenty-two wheat powdery mildew samples were collected in 2018 from four locations in the Tasmanian broadacre cropping region during a two-day collection trip (Figure 1 and Table 1). Samples were maintained in polypropylene tubes containing benzimidazole agar, as previously described, and stored in a polystyrene box with ice bricks to prevent deterioration prior to processing.

**Figure 1.**
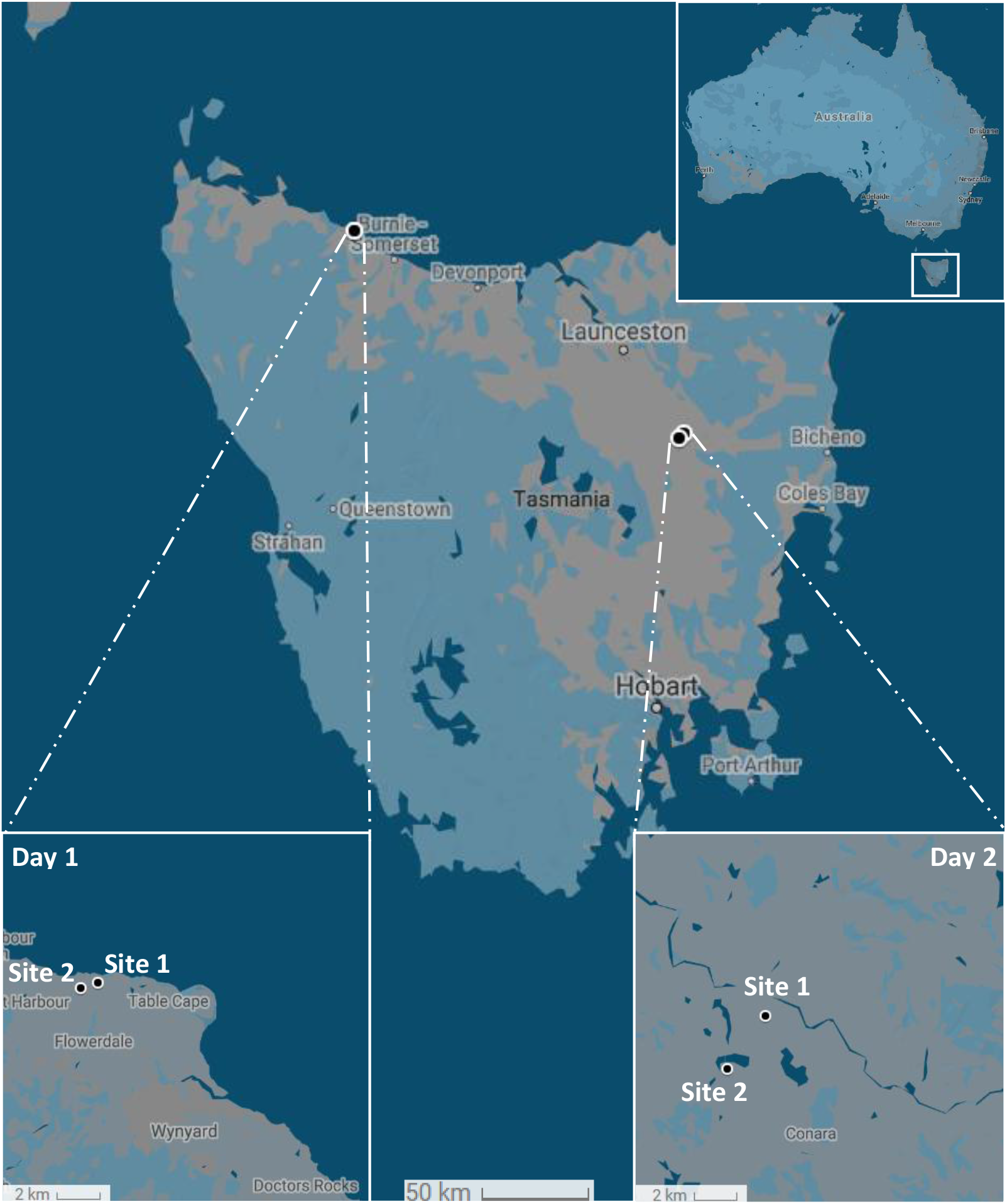
Map showing sampling locations in Tasmania (Australia). Sampling on day 1 was conducted in Tasmania’s Cradle-coast/North-western region, near Table Cape. Sampling on day 2 was conducted in the Northern/Northern Midlands region, near Conara.

On each day, once sampling was concluded, sixteen 7 mm discs per infected leaf sample were excised using a WellTech Rapid-Core-7.0 (Rapid Core, Australia) disinfected with 70% v/v ethanol between samples. The discs were halved and separated into two 1.5 ml microfuge tubes to obtain two sets of tubes (1 and 2) with 16 halves each, per sample. The material in set 1 was used for immediate processing and that of set 2 shipped on ice by overnight courier to our laboratory in Perth for dPCR analysis (Figure 2). Samples in set 1 were processed using the quick DNA extraction method described earlier. The ASqPCR reactions were set-up as described above and 5 μl of the DNA extract was used directly in the reaction mix. Each sample was tested in triplicate and amount of fungal DNA was measured using a standard curve constructed from Goomalling and Geelong 1 DNA of known concentrations amplified alongside the samples. The amount of mutant allele in each sample was reported as a percent value. Remaining crude DNA was shipped with the tubes from set 2 for subsequent dPCR analysis in the laboratory.

**Figure 2.**
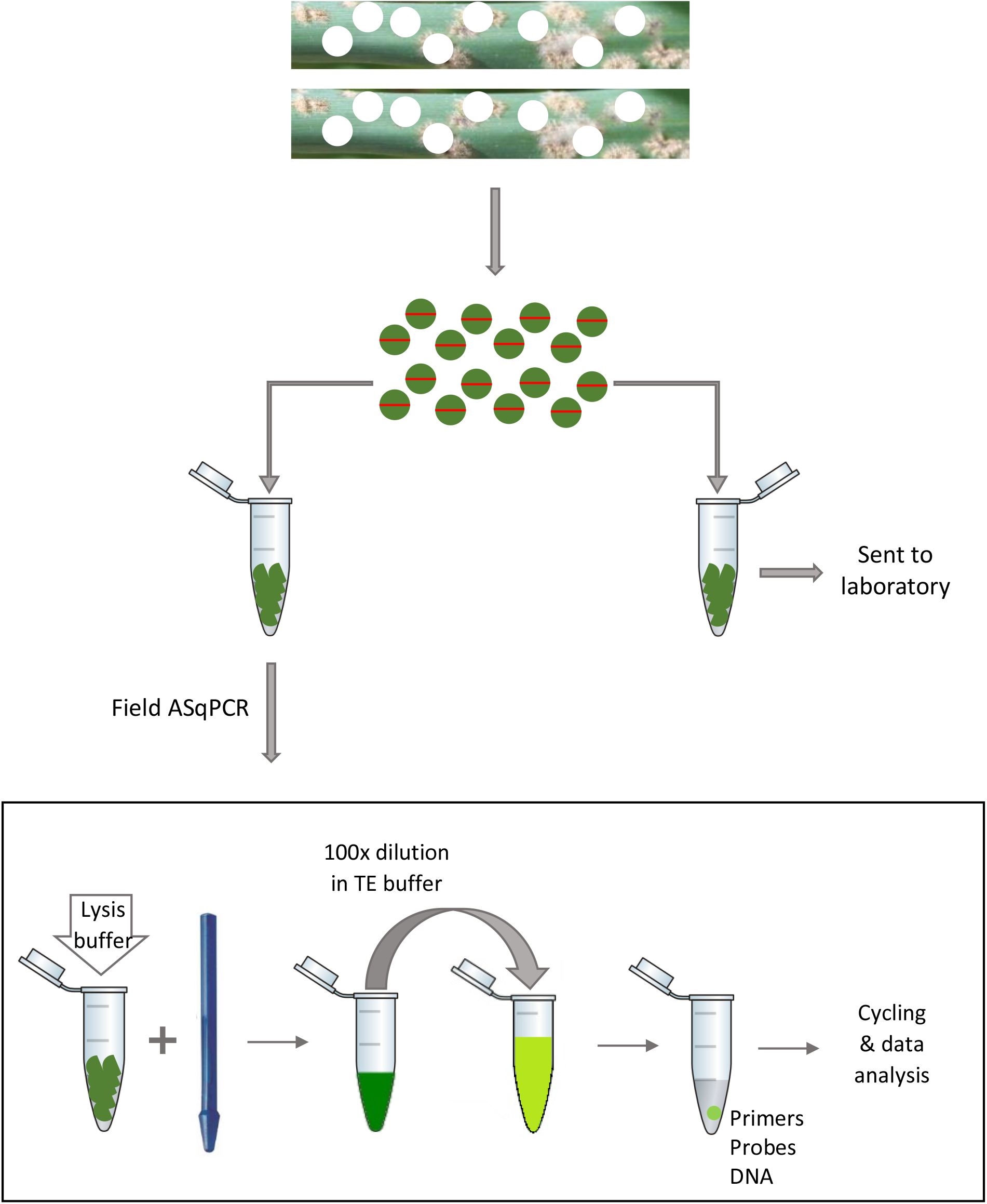
Schematic representation of the procedure followed in the field for the extraction of DNA.

### In lab sample processing

Upon reception of set 2 in the laboratory, tissue was crushed to a fine powder using a 2 mm stainless steel ball bearing in a Retsch mixer mill MM400 (Retsch, Germany), followed by DNA extraction with a Biosprint Plant DNA 15 kit (Qiagen), as per manufacturer’s instructions. The DNA was eluted in 200 μl of 1x TE buffer and tested using both dPCR and ASqPCR in the same manner as above. Any sample that fell outside the 200-2,000 copy range on the dPCR was diluted/concentrated and re-tested as required.

### Statistical analysis of dPCR and ASqPCR results

The results of the dPCR and ASqPCR analyses were compared using SigmaPlot v13 (Systat Software, San Jose, CA). A Bland-Altman analysis, also known as the Tukey’s mean difference test, and a paired samples t-test were conducted. The Bland-Altman plot was constructed by plotting the difference between the paired data points (method 1 *minus* method 2) on the y-axis and the mean of the paired data points on the x-axis.^30^ The mean of the difference between the pairs represents the bias i.e. whether the method 1 is overestimating (positive bias) or underestimating (negative bias) compared to method 2. The upper and lower limits of confidence are calculated as the mean (bias) +/− 1.96 standard deviation of the differences.

### In-field application of the ASqPCR pipeline for the detection of cytb mutation g428c

In October 2019, suspected strobilurin-resistant wheat powdery mildew samples (n=12) brought by growers attending an industry field day at Northern Yorke Peninsula (South Australia), were processed in a field (Table 1). The tissue was not excised using the coring tool, rather cut into small fragments using scissors and forceps disinfected with 70% v/v ethanol between samples. Each sample was added up to the 100 μl graduation in 1.5 ml microfuge tubes. This was followed by a quick DNA extraction and ASqPCR assay on the MIC instrument as detailed above, and mutant frequency results communicated to the growers upon assay completion.

## Results

### Identification of the G143A mutation in the CytB protein

A 465bp PCR amplicon of the *Bgt cytb* gene was amplified from 10 wheat powdery mildew and one barley powdery mildew samples and sequenced as described above (Tables 1 and 2). The alignment of these sequences together with QoI-sensitive and resistant *Bgt cytb* reference sequences, revealed the presence of the mutation *g428c* (Cytb amino acid substitution G143A) in two samples each from Tasmania and Victoria collected from paddocks where the use of QoI fungicides failed to control the disease (Figure 3). The analysis of the chromatograms also show that for the Victorian samples Geelong 1 and 2 there was only a peak for a C base at the mutation site, where as for the Tasmanian samples Launceston 1 and 2, there was an additional smaller peak for a G base. All samples analysed from Western Australia and South Australia were wild-type.

**Figure 3.**
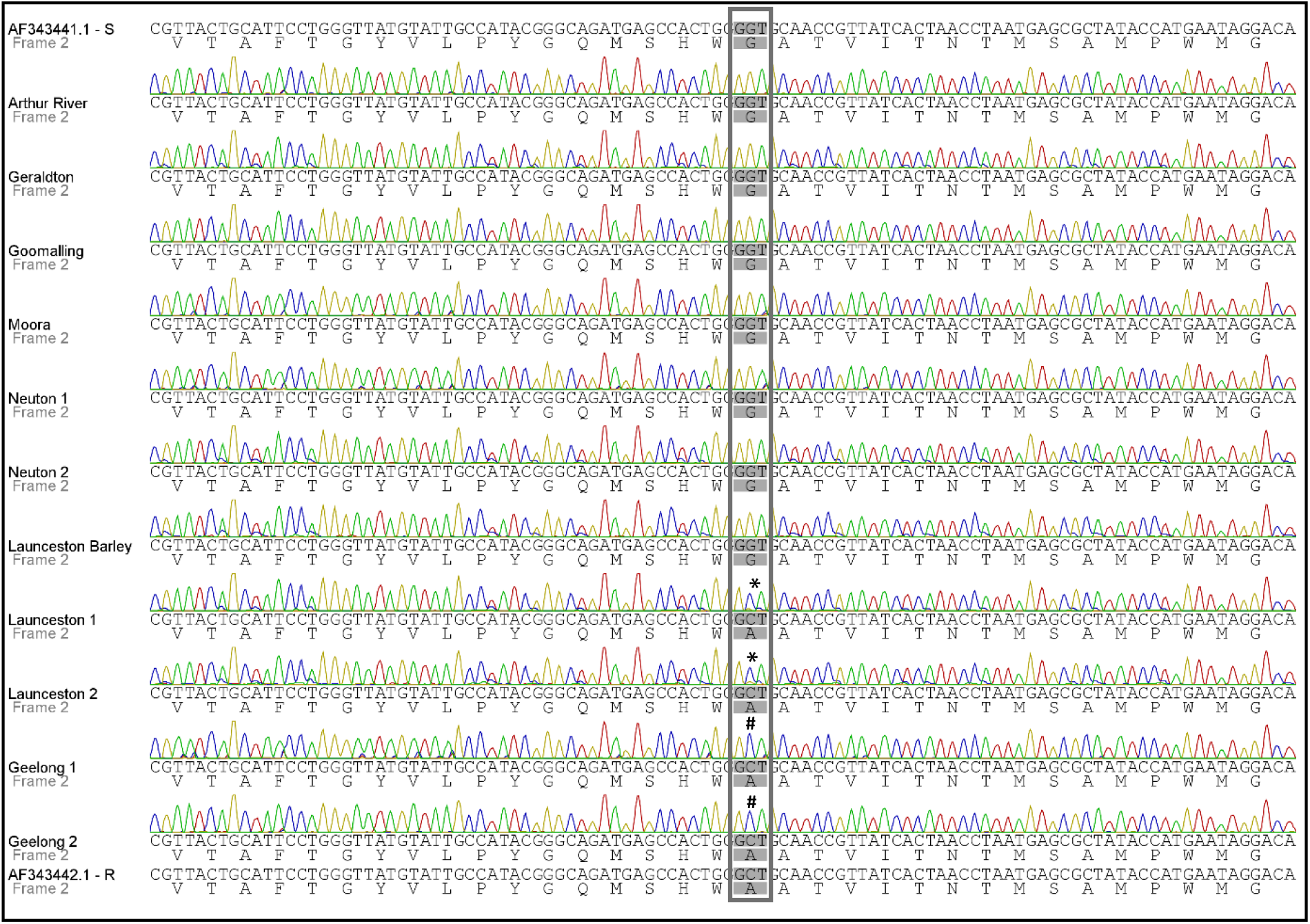
Multiple sequence alignment of the *cytb* gene sequence and its translation (Yeast mitochondrial translation table_3) flanking the *c. G>C 428* mutation for samples collected in 2016 (Table 1). AF343441.1 and AF343442.1 correspond to strobilurin sensitive and resistant *Bgt cytb* reference sequences, respectively. The samples Launceston 1 and 2, and Geelong 1 and 2 contain the substitution G143A. Codon 143 is boxed; * denotes a mix of the G and C base; # shows C base only.

### dPCR and ASqPCR successfully quantify Bgt cytb g428c

The dPCR analysis successfully detected the A143 allele in total DNA from field samples collected in 2016 corroborating the results from the sequencing of the *cytb* gene (Tables 1 and 3; Figure 3). To evaluate the accuracy and sensitivity of the assay, genomic DNA from QoI-sensitive and resistant disease samples Goomalling (G143) and Geelong 1 (A143), was mixed in known ratios and subjected to dPCR analysis. Scatterplot representation of the results showed well-defined, discrete groups of either wild type or mutant allele only, both alleles and neither allele present (Figure 4A). The analysis of the dilution series indicated a good correlation between percentage of mutant detected and known mutant/wild type ratios (R^2^= *0.999*), with the lower level of quantification of the mutant allele determined to be 0.33%, with the allele detectable at 0.07% (Figure 4B, Table 4). No cross-reaction was found when genomic DNA from other common wheat pathogens was tested, however, the closely related barley powdery mildew pathogen *B. graminis* f. sp. *hordei* was positive for the G143 allele (Table 3).

**Table 3.**
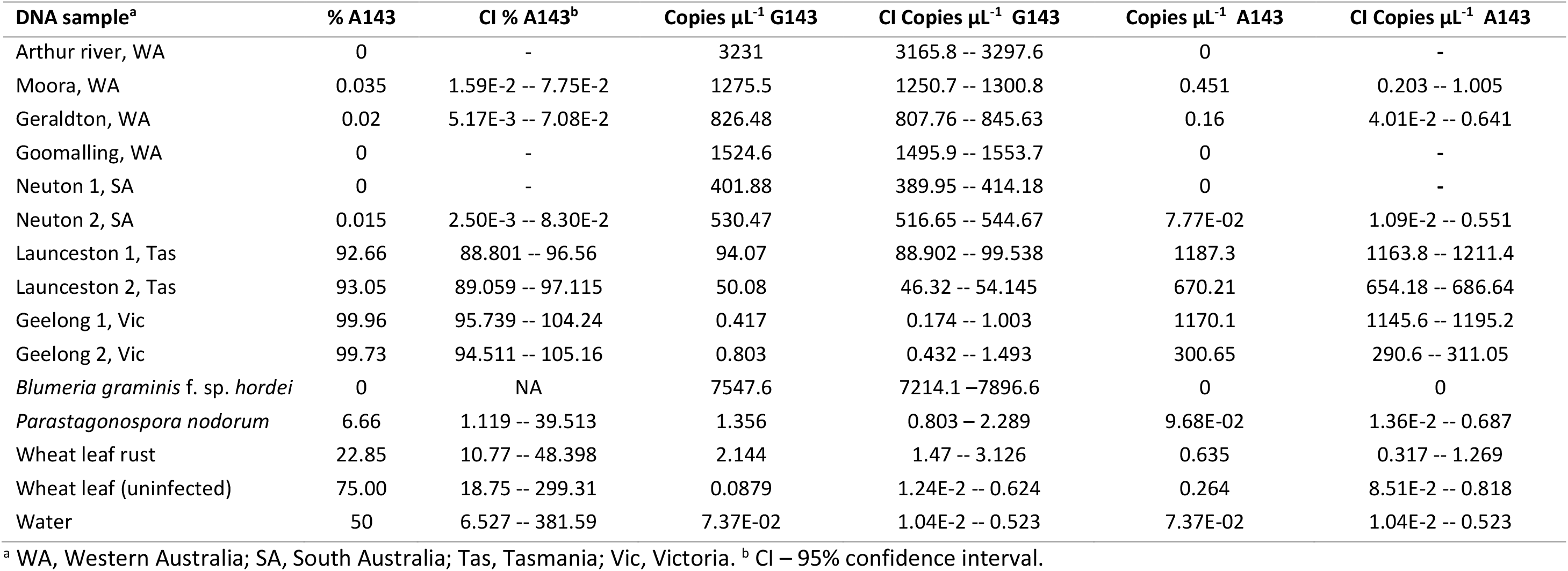
Specificity of detection of the G143A dPCR assay on total DNA from *Blumeria graminis* f. sp. *tritici* field samples and controls.

| DNA sample<sup>a</sup> | % A143 | CI % A143<sup>b</sup> | Copies $\mu\text{L}^{-1}$ G143 | CI Copies $\mu\text{L}^{-1}$ G143 | Copies $\mu\text{L}^{-1}$ A143 | CI Copies $\mu\text{L}^{-1}$ A143 |
| --- | --- | --- | --- | --- | --- | --- |
| Arthur river, WA | 0 | - | 3231 | 3165.8 -- 3297.6 | 0 | - |
| Moora, WA | 0.035 | 1.59E-2 -- 7.75E-2 | 1275.5 | 1250.7 -- 1300.8 | 0.451 | 0.203 -- 1.005 |
| Geraldton, WA | 0.02 | 5.17E-3 -- 7.08E-2 | 826.48 | 807.76 -- 845.63 | 0.16 | 4.01E-2 -- 0.641 |
| Goomalling, WA | 0 | - | 1524.6 | 1495.9 -- 1553.7 | 0 | - |
| Neuton 1, SA | 0 | - | 401.88 | 389.95 -- 414.18 | 0 | - |
| Neuton 2, SA | 0.015 | 2.50E-3 -- 8.30E-2 | 530.47 | 516.65 -- 544.67 | 7.77E-02 | 1.09E-2 -- 0.551 |
| Launceston 1, Tas | 92.66 | 88.801 -- 96.56 | 94.07 | 88.902 -- 99.538 | 1187.3 | 1163.8 -- 1211.4 |
| Launceston 2, Tas | 93.05 | 89.059 -- 97.115 | 50.08 | 46.32 -- 54.145 | 670.21 | 654.18 -- 686.64 |
| Geelong 1, Vic | 99.96 | 95.739 -- 104.24 | 0.417 | 0.174 -- 1.003 | 1170.1 | 1145.6 -- 1195.2 |
| Geelong 2, Vic | 99.73 | 94.511 -- 105.16 | 0.803 | 0.432 -- 1.493 | 300.65 | 290.6 -- 311.05 |
| *Blumeria graminis* f. sp. *hordei* | 0 | NA | 7547.6 | 7214.1 -- 7896.6 | 0 | 0 |
| *Parastagonospora nodorum* | 6.66 | 1.119 -- 39.513 | 1.356 | 0.803 -- 2.289 | 9.68E-02 | 1.36E-2 -- 0.687 |
| Wheat leaf rust | 22.85 | 10.77 -- 48.398 | 2.144 | 1.47 -- 3.126 | 0.635 | 0.317 -- 1.269 |
| Wheat leaf (uninfected) | 75.00 | 18.75 -- 299.31 | 0.0879 | 1.24E-2 -- 0.624 | 0.264 | 8.51E-2 -- 0.818 |
| Water | 50 | 6.527 -- 381.59 | 7.37E-02 | 1.04E-2 -- 0.523 | 7.37E-02 | 1.04E-2 -- 0.523 |

<sup>a</sup> WA, Western Australia; SA, South Australia; Tas, Tasmania; Vic, Victoria. <sup>b</sup> CI – 95% confidence interval.

**Figure 4.**
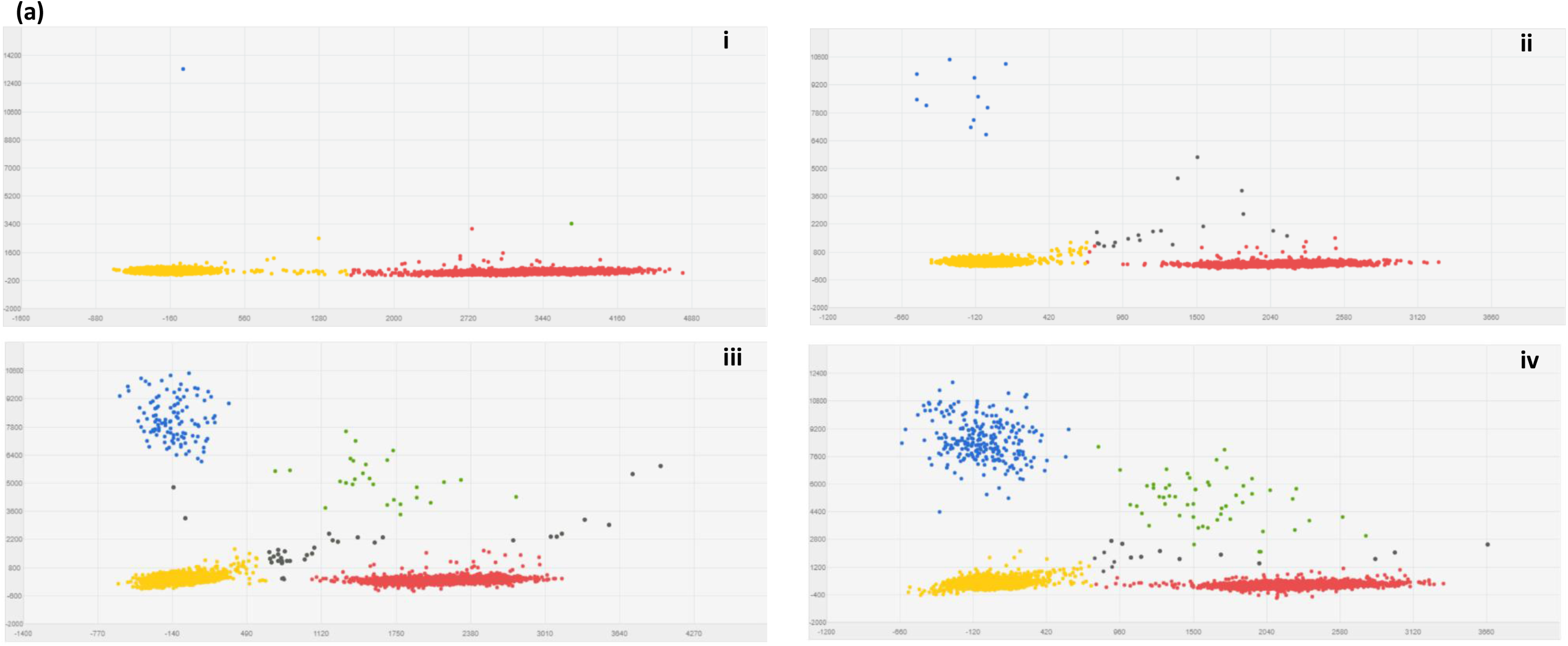
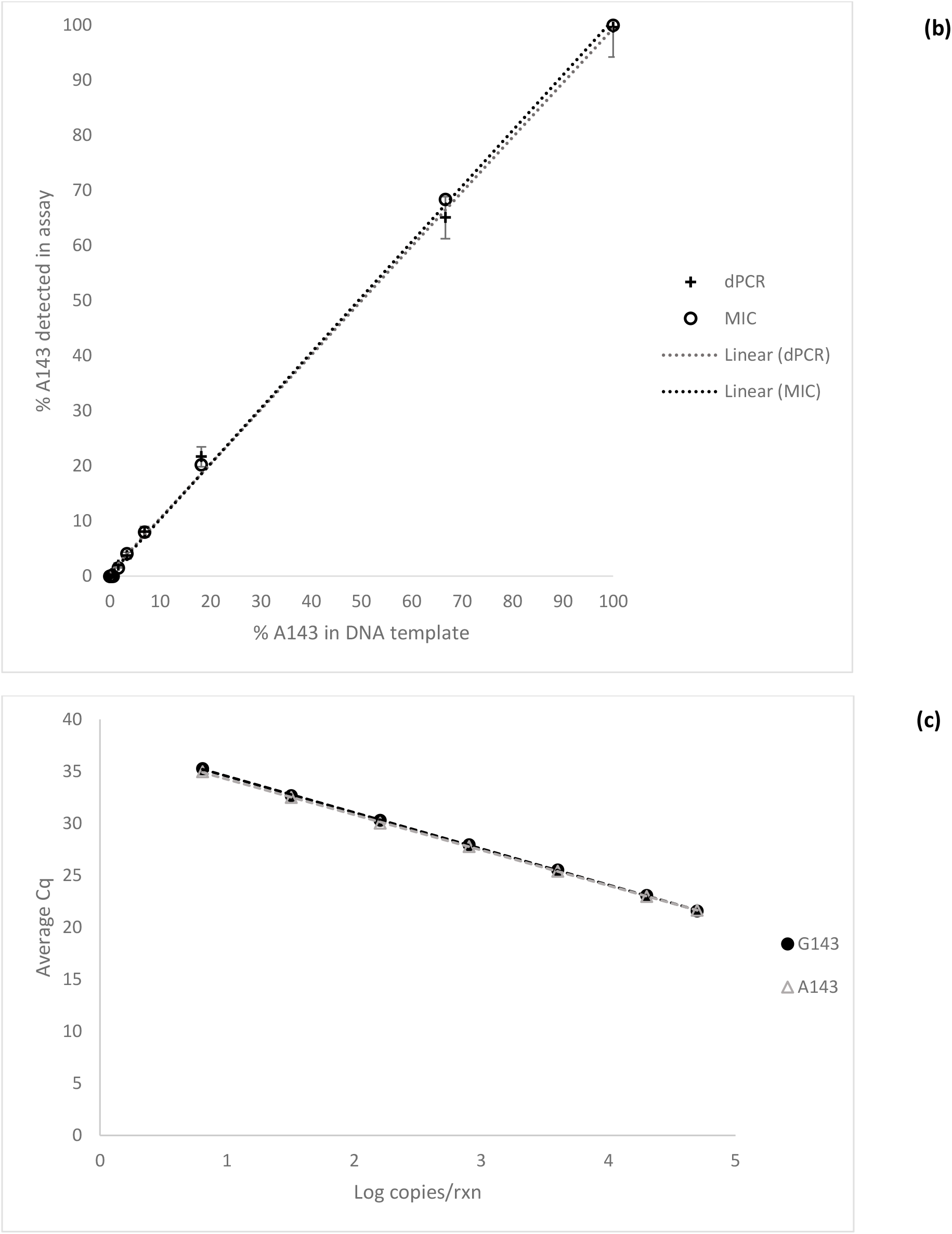
Sensitivity and specificity of the dPCR and ASqPCR assays targeting the *cytb* G143A. (a) Scatter plots for *Blumeria graminis* f. sp *tritici* (*Bgt*) G143A dPCR assay using known ratios of genomic DNA for the strobilurin resistant *Bgt* sample Geelong 1 (A143 allele) and sensitive *Bgt* sample Goomalling (G143 allele). Scatter plots were prepared with Quantstudio™ 3D AnalysisSuite™. Wells with the A143 allele are represented by blue signals (FAM), G143 alleles are represented by red signals (VIC), detection of both alleles are represented by green signals, wells without any alleles (passive reference) are represented by yellow signals (ROX) and outliers represented by grey signals. (i) 100% Goomalling gDNA (G143 allele only); A143 frequency = 0.04%. (ii) 0.33% Geelong1 gDNA; A143 frequency = 0.24%. (iii) 3.39% Geelong 1 gDNA; A143 frequency = 3.10%. (iv) 6.90% Geelong 1 gDNA; A143 frequency = 7.87%. (b) Linear correlation between percent G143A mutant *Blumeria graminis* f. sp. *tritici* (A143) genomic DNA samples quantified by digital PCR (R^2^ = 0.9988) and ASqPCR (R^2^ = 0.9994), and the input percentage of the mutant in a genomic DNA mixture of A143 and G143. Each point represents the average of triplicates. (c) Correlation of threshold cycle in ASqPCR and amount of DNA of either G143 (R^2^ = 0.9998) or A143 (R^2^ = 0.9999) alleles, respectively.

**Table 4.**
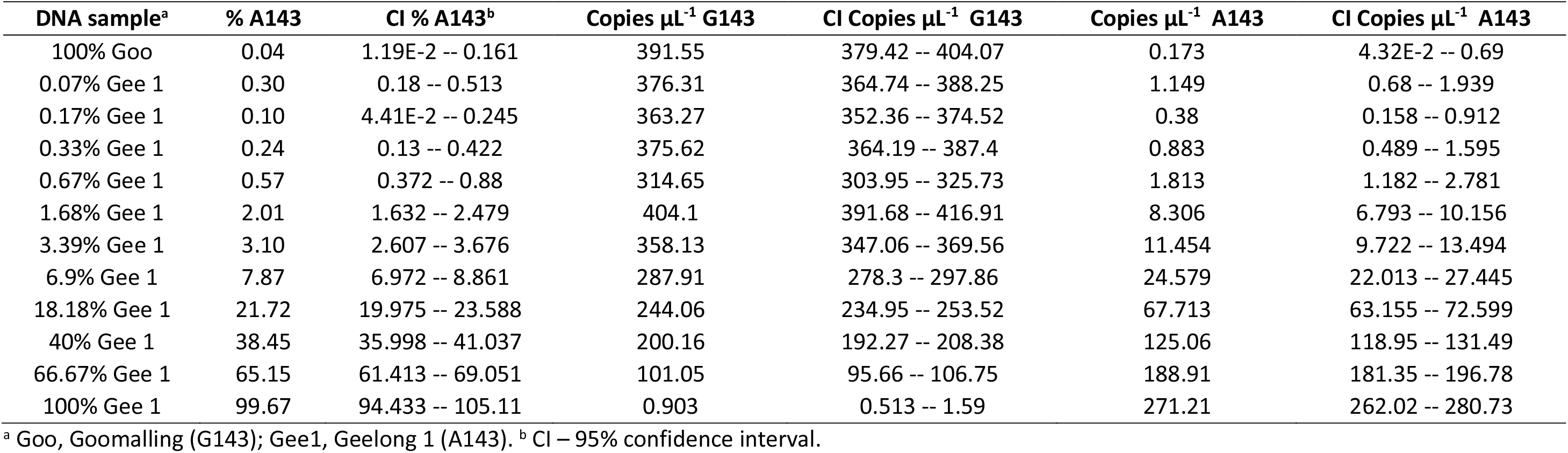
Sensitivity of the G143A dPCR assay on a mixture of total DNA from mutant (A143) and wild type (G143) *Blumeria graminis* f. sp. *tritici* field samples.

| DNA sample<sup>a</sup> | % A143 | CI % A143<sup>b</sup> | Copies $\mu\text{L}^{-1}$ G143 | CI Copies $\mu\text{L}^{-1}$ G143 | Copies $\mu\text{L}^{-1}$ A143 | CI Copies $\mu\text{L}^{-1}$ A143 |
| --- | --- | --- | --- | --- | --- | --- |
| 100% Goo | 0.04 | 1.19E-2 -- 0.161 | 391.55 | 379.42 -- 404.07 | 0.173 | 4.32E-2 -- 0.69 |
| 0.07% Gee 1 | 0.30 | 0.18 -- 0.513 | 376.31 | 364.74 -- 388.25 | 1.149 | 0.68 -- 1.939 |
| 0.17% Gee 1 | 0.10 | 4.41E-2 -- 0.245 | 363.27 | 352.36 -- 374.52 | 0.38 | 0.158 -- 0.912 |
| 0.33% Gee 1 | 0.24 | 0.13 -- 0.422 | 375.62 | 364.19 -- 387.4 | 0.883 | 0.489 -- 1.595 |
| 0.67% Gee 1 | 0.57 | 0.372 -- 0.88 | 314.65 | 303.95 -- 325.73 | 1.813 | 1.182 -- 2.781 |
| 1.68% Gee 1 | 2.01 | 1.632 -- 2.479 | 404.1 | 391.68 -- 416.91 | 8.306 | 6.793 -- 10.156 |
| 3.39% Gee 1 | 3.10 | 2.607 -- 3.676 | 358.13 | 347.06 -- 369.56 | 11.454 | 9.722 -- 13.494 |
| 6.9% Gee 1 | 7.87 | 6.972 -- 8.861 | 287.91 | 278.3 -- 297.86 | 24.579 | 22.013 -- 27.445 |
| 18.18% Gee 1 | 21.72 | 19.975 -- 23.588 | 244.06 | 234.95 -- 253.52 | 67.713 | 63.155 -- 72.599 |
| 40% Gee 1 | 38.45 | 35.998 -- 41.037 | 200.16 | 192.27 -- 208.38 | 125.06 | 118.95 -- 131.49 |
| 66.67% Gee 1 | 65.15 | 61.413 -- 69.051 | 101.05 | 95.66 -- 106.75 | 188.91 | 181.35 -- 196.78 |
| 100% Gee 1 | 99.67 | 94.433 -- 105.11 | 0.903 | 0.513 -- 1.59 | 271.21 | 262.02 -- 280.73 |

<sup>a</sup> Goo, Goomalling (G143); Gee1, Geelong 1 (A143). <sup>b</sup> CI – 95% confidence interval.

The assay identified very high frequency levels of the A143 allele in locations where QoI control failure had previously been reported (Tables 1 and 3). Where QoIs controlled the disease satisfactorily, the frequency of the A143 allele remained below the assay detection level. The double peaks observed in the sequencing chromatograms of samples Launceston 1 and 2 were in agreement with the A143 allele frequencies determined by the dPCR assay (92.66 and 93.05%, respectively). In a similar manner, samples Geelong 1 and 2 showed A143 frequencies of virtually 100%, which correlated with the single peaks observed in the chromatograms of these two samples (Figure 3 and Table 3).

The SensiFast Probe No-Rox mastermix was considered the most suitable for ASqPCR with crude DNA samples on the MIC instrument under field conditions. In the reactions where the iQ multiplex Supermix was used, the G143 allele was detected in both the HEX and FAM channels. In the case of the samples analysed using the Immomix™, no amplification was observed from any of the crude DNA extracts (Table 5).

**Table 5.** Cq values obtained from the assessment of three qPCR mastermixes.

|  | G143 target (HEX channel) |  |  | A143 target (FAM channel) |  |  |
| --- | --- | --- | --- | --- | --- | --- |
|  | Mix 1 | Mix 2 | Mix 3 | Mix 1 | Mix 2 | Mix 3 |
| 100% G143 (L)<sup>a</sup> | 19.02 ±0.50<sup>d</sup> | 19.22 ±0.50 | 34.25 ±1.21 | **25.65 ±0.34** | - | - |
| 100% G143 (Q)<sup>b</sup> | 23.79 ±0.56 | 24.49 ±0.44 | - | - | - | - |
| 100% A143 (L) | - | - | - | 18.39 ±0.40 | 18.64 ±0.35 | 32.52 ±0.22 |
| 100% A143 (Q) | - | - | - | 23.43 ±0.25 | 23.89 ±0.22 | - |
| 51% A143 (L)<sup>c</sup> | 23.53 ±0.63 | 23.78 ±0.65 | 38.24 ±0.09 | 21.60 ±0.09 | 21.99 ±0.15 | 36.73 ±0.27 |
| NTC | - | - | - | - | - | - |

Mix 1 (iQ Multiplex Supermix), Mix 2 (Sensifast Probe No-Rox mix) and Mix 3 (Immomix™), with total DNA extracted from wheat powdery mildew wild type (G143) and mutant (A143) field samples. **Non-target amplification is indicated in bold.** <sup>a</sup> DNA extracted using a standard laboratory method. <sup>b</sup> DNA extracted using the quick field extraction method. <sup>c</sup> 49% G143. <sup>d</sup> Standard deviation (5 replicates)

The lower level of quantification of the mutant allele in mixture with the wild type was in this case 1.67% (Figure 4B). In addition, in homoplasmic reactions, the assay quantified down to seven copies per reaction for both G143 and the A143 alleles (Figure 4C). As with the dPCR assay, wheat pathogens *P. nodorum* and *P. tritici-repentis* were not detected. The assay however detected the closely related barley mildew *B. graminis* f.sp. *hordei*, but not the grape mildew *E. necator* (data not shown).

### Optimisation of the in-field allele frequency quantification pipeline

The suitability of the in-field *Bgt cytb* A134 allele quantification method was assessed as part of a two-day field trip across the Northern wheat-growing region of Tasmania (Figure 1). The ASqPCR assay conducted on the MIC instrument, with crude DNA extracted in the field as described above, successfully distinguished between G143 and A143 alleles under field conditions. The resistant genotype could be detected in all of the 22 samples tested across the four collection sites, albeit at varying levels (Figure 5).

**Figure 5.**
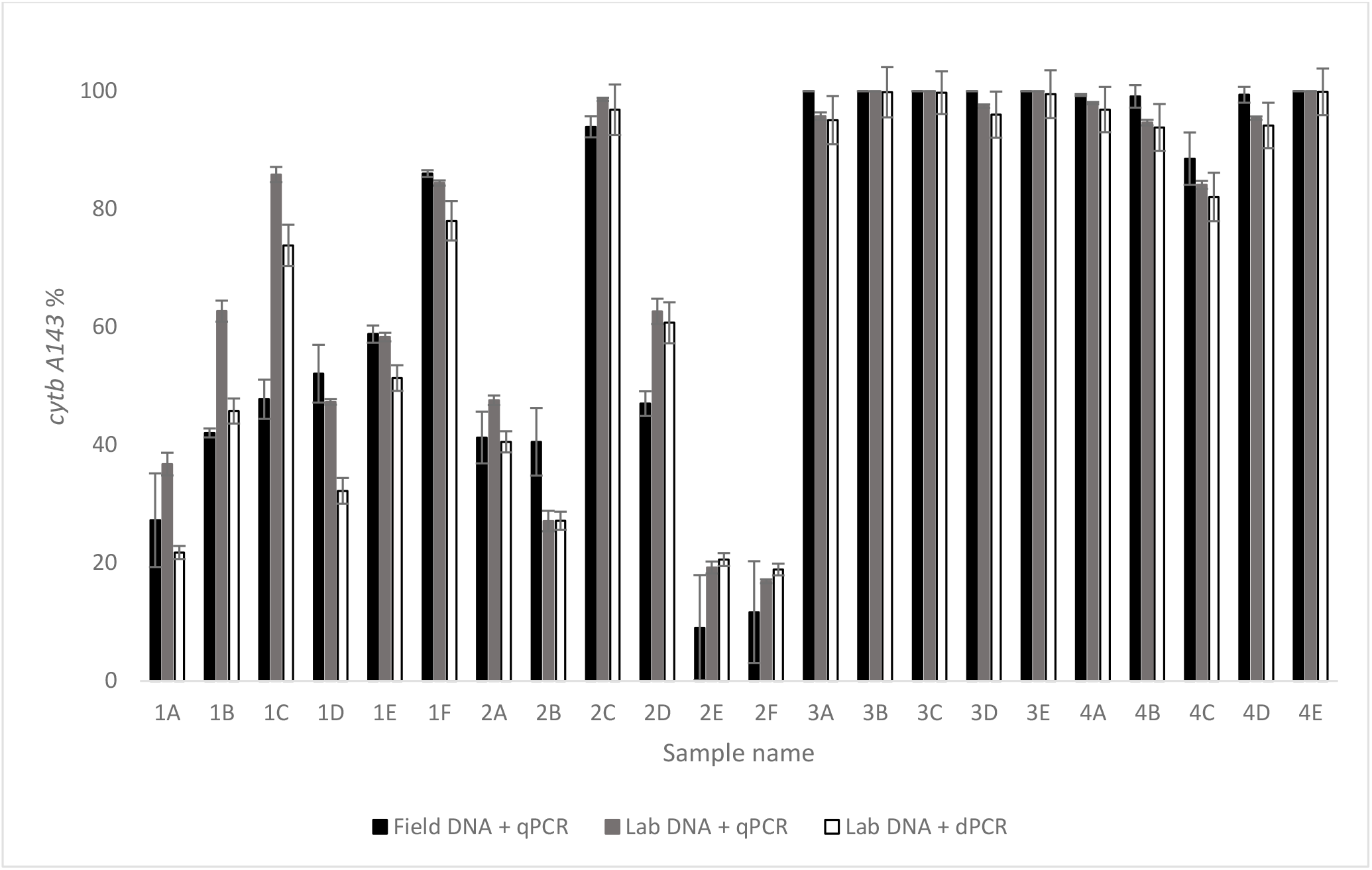
Comparison of *cytb* A143 frequency results of samples analysed using a field qPCR (Field DNA + qPCR), laboratory qPCR (Lab DNA + qPCR) and laboratory dPCR (Lab DNA + dPCR). 1A-1F and 2A-2F –samples collected from sites 1 and 2 in the Table Cape region; 3A-3E and 4A-4E –samples collected from sites 1 and 2 in the Conara region. Error bars show standard error of mean.

The largest variation in the frequency of the *cytb* mutant allele was observed in the samples collected on day 1. Frequencies of the A143 allele in the samples from Table Cape ranged from 27.26% - 86.04% (average = 52.34% ±19.67) in the first site (Figure 5, 1A-1F) and from 9.03 - 93.98% (average = 40.60 ±30.74) in the second site (Figure 5, 2A-2F). The analysis of the samples collected at the two sites in the Conara revealed a higher frequency of the A143 allele, with the majority of the samples showing values above 99% (average = 98.65 ±3.56) (Figure 5, 3A-3E and 4A - 4E).

### In-lab analysis of samples

The second set of powdery mildew infected wheat disc halves was sent by overnight courier and analysed in our laboratory as described earlier. The frequency of the A143 allele in both the dPCR and ASqPCR analyses was comparable to the results from the field analysis (Figure 5). Samples collected on day 1 had, in general, lower and more variable A143 allele frequencies than those from day 2. The frequency of the mutant allele in the samples collected from site one was 36.78% to 85.89% and 21.78% to 78.03% for the dPCR and ASqPCR assays, respectively (Figure 5, 1A - 1F). For these analyses as well, the largest variation in the A143 frequency was found in the samples from the second site, which had frequencies ranging from 16.90% to 98.64% for dPCR and 18.89% to 96.83% for ASqPCR (Figure 5, 2A - 2F).

As with the in-field analysis, the mutant allele was found at frequencies close to 100% in the majority of samples collected on day 2. In this case, the frequency of the A143 allele observed ranged from 82.03% to 99.92% and 84.11 to 100% for dPCR and ASqPCR, respectively (Figure 5, 3A – 4E). No results were obtained when the crude extract which had been shipped with the set 2 of infected material, was analysed using the dPCR assay (data not shown).

### Comparison of test methods

To determine the accuracy of the in-field ASqPCR method, A143 allele frequency data was compared to that from ASqPCR and dPCR experiments performed in our laboratory. Paired sample t-test analysis showed that the frequency results obtained in the field were not significantly different from those of the same experiment performed in the laboratory using a standard method to extract DNA from the original samples (*p* value = 0.190). The same was observed when the A143 allele frequencies from the field analysis were compared to the results obtained with dPCR using DNA extracted in the laboratory (*p* value = 0.677).

The Bland-Altmann analysis also showed a good correlation between the field ASqPCR and the laboratory dPCR methods, with data points distributed about the line of bias (Figure 6A). On average, the A143 allele frequencies calculated using the field ASqPCR were overestimated by 0.86% (bias), compared to the dPCR results. Twenty out of 22 samples (90.91%) were within the upper and lower limits of agreement of 19.21 and −17.48, respectively (Figure 6A).

**Figure 6.**
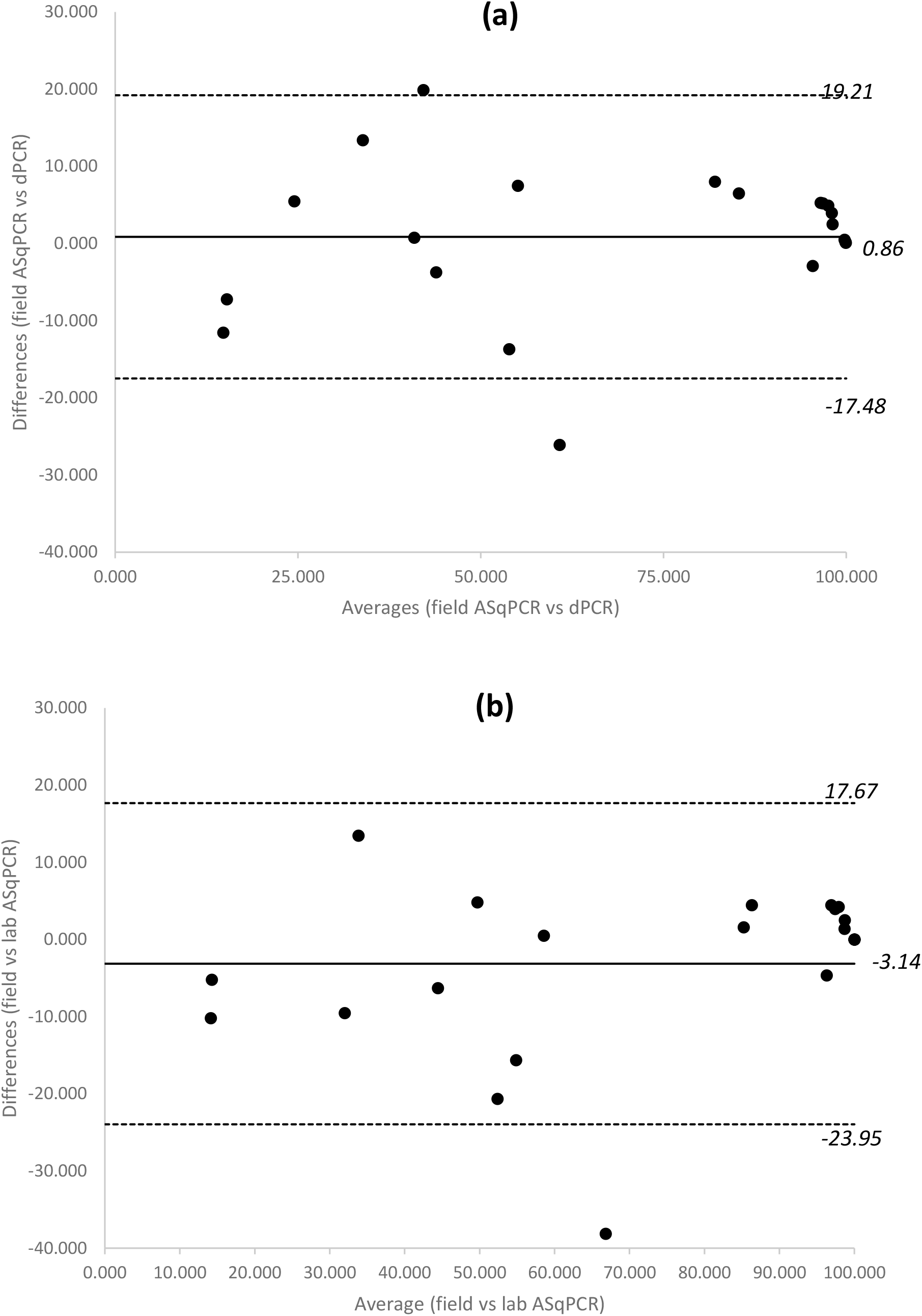
Bland-Altman analysis to check agreement between the results from the field qPCR and lab dPCR or lab ASqPCR (n=22). (a) Plot of the differences and means for the results from ASqPCR and dPCR; the bias of 0.86 represented the average difference between the results from the methods and shows that the results from the ASqPCR are overestimated by 0.86 compared to the latter. The upper and lower limits of agreements at 19.21 and −17.48 respectively represent the bias ±1.96 SD (SD of differences is 9.36) 90.1% of samples are within the limits of agreement. (b) Plot of the differences and means for the results from field ASqPCR and lab ASqPCR; the bias of −3.14 represents the average difference between the results from the methods and shows that the results from the crude DNA are underestimated by 3.14 compared to the latter. The upper and lower limits of agreements at 17.67 and −23.95 respectively represent the bias ±1.96 SD (SD of differences is 10.62) 95.5% of samples are within the limits of agreement.

The same was true for the ASqPCR conducted with either the crude or laboratory DNA extract, with data points distributed about the line of bias (Figure 6B). In this case, the allele frequencies calculated using the field ASqPCR are underestimated by 3.14%, compared to the results from the laboratory DNA. Twenty-one out of 22 samples (95.5%) were within the upper and lower limits of agreement of 17.67 and −23.95, respectively (Figure 6B).

### Deployment of the field Bgt cytb G143A ASqPCR

Towards the end of the 2019 growing season, South Australian agronomists were reporting poor control of wheat powdery mildew after treatment with QoI fungicides. The A143 ASqPCR method was showcased as part of an extension activity organised within an industry field day in Bute, Yorke Peninsula (South Australia). A total of 12 symptomatic samples were brought by participants from surrounding wheat fields and analysed on the spot. The frequency of the *cytb* A143 allele ranged from 1.94 - 53.72% (average = 14.19 ±17.41) suggesting a relatively high spread of the mutation in the area (Table 1). Three samples had no detectable levels of the A143 allele.

## Discussion

Failure of QoI (quinone outside inhibitor) fungicides to control *Blumeria graminis* f. sp. *tritici* (*Bgt*) in wheat paddocks in the Australian states of Tasmania and Victoria was first reported in 2016 (F. Lopez-Ruiz, personal communication). Sequencing analysis of the *cytb* gene revealed that the well characterised mutation G143A was present in all samples collected from wheat fields in these two Australian states where adequate control of the disease could not be achieved with the use of QoI fungicides (Figure 3).^21^ As the G143A genotype continues to spread, and possibly emerge, across other wheat growing regions in the world, due to the overexposure to this group of fungicides, the availability of a rapid and robust field-based allele specific detection method could provide a key advantage for the in-season management of QoI resistance.

The detection of fungicide resistance in crop pathogens, previously mostly conducted using bioassays, now largely relies on the use of molecular methods when the genetic mechanism associated with the resistance is known.^31^ For novel field failures, where the target gene is known, capillary sequencing methods are used to determine if any known mutations are the cause.^32^ Once a mutation has been confirmed, PCR and LAMP are commonly used for its detection, whereas for quantification qPCR and more recently digital PCR are used.^12,17,29,33,34^ Except for LAMP, none of the other methods have been used in the field for the detection of fungicide resistance, therefore currently the quantification of mutations associated with fungicide resistance is exclusively laboratory based.^14^ We have demonstrated the successful development of a TaqMan probe based ASqPCR assay, which allows for both the mutant and wild-type alleles to be detected within the same reaction. This has been coupled with a quick, field-friendly DNA extraction method and a robust DNA polymerase to enable the in-field quantification of the QoI resistant mutation G143A in *Bgt* on a lightweight and fast qPCR instrument.

The quantification of the CytB G143A mutation in *Bgt* using ASqPCR was first conducted by Fraaije et al.^17^ using the intercalating dye SYBR 1, with the assay detecting as low as 1 in 10,000 copies of the mutant A134 allele in a sample extraction that contained a mix of fungal and wheat DNA. A qPCR assay developed for the analysis of mutation Y136F in the Cyp51 target site of *E. necator* had a limit of detection and a limit of quantification of 0.85 and 2.85%, respectively.^35^ More recently, Zulak et al.^29^ developed a chip dPCR assay able to quantify mutations S509T and Y136F in the Cyp51 of *B. graminis* f.sp. *hordei* infected samples down to 0.2%. In our study, the quantification limit of the field-based G143A ASqPCR assay when using a DNA extract containing a mixture of both A143 and G143 alleles, and wheat DNA, was 1.67% (Figure 4B). However, in samples with homogenous (homoplasmic) genotypes, in a background of wheat DNA, the assay successfully detected seven copies of the G143 or A143 alleles per reaction (Figure 4C). This could be considered as the equivalent to a single spore detection, as each spore has multiple mitochondria and each mitochondrion may have multiple copies of the *cytb* gene within it.^36^ A laboratory based G143A chip dPCR assay was also developed for comparative purposes. The G143A dPCR assay successfully quantified the *cytb* A143 allele in a background of G143 and wheat DNA down to a level of 0.33%, whereas its detection limit was estimated at 0.07% (Figures 4A and B; Table 4). dPCR has, in general, lower target detection limits and higher reproducibility than qPCR, the latter due to the use of absolute quantification rather than relying on standard curves.^29,37,38^ Due to these properties, dPCR could be considered the gold-standard in molecular detection of fungicide resistant populations.

The assessment of the ASqPCR analysis pipeline under field conditions revealed a marked A143 allele frequency difference between two wheat growing regions of Tasmania (Figure 5). While the average frequencies of the A143 allele in samples collected at two sites in Table Cape remained between 40-50%, the values found in the sites sampled in Conara showed levels above 99% in nine out of ten analysed samples. The accuracy of the in-field G143A ASqPCR method was validated by comparing the A143 allele frequency results obtained in the field to those from the laboratory using the same method with DNA extracted from the original samples following standard laboratory procedures. The lack of significant differences between the two datasets indicated that neither the quick in-field DNA extraction nor the lack of a controlled laboratory environment had an important effect on the outcome of the in-field analysis (*p* value=0.190).

To assess the level of sensitivity of the in-field G143A ASqPCR method, we compared the A143 allele frequency results to those obtained from dPCR using Bland-Altman analysis. When this analysis is applied to two testing methods and the results are identical, the average of the differences between paired results should be zero.^30^ In this study, the comparison of the two tests indicated that ASqPCR is, on average, overestimating the mutant allele in a sample by just 0.86 units (line of bias; Figure 5). This is quite remarkable considering that the dPCR method used DNA extracted in the laboratory as opposed to the crude extractions analysed in the field. In our analysis, the limits of agreement, which ranged from 19.21 to −17.48 units, represents 95% of the range of the differences between the measurements, and eight of the 12 differences greater than 5% are due to the field analysis showing higher frequencies than the dPCR analysis. The biggest differences were observed in the samples from the Table Cape region where the more moderate A134 levels seemed to display better the overestimating effect (Figure 5, 1A - 2F). This effect, which is likely to be a consequence of the higher sensitivity of the dPCR assay, can also be influenced by the fact that the DNA analysed by the two methods was extracted from different areas of the same leaf disc.

The fact that the Northern region has almost double the area dedicated to broadacre cropping (60,278 ha) than the North Western region (31,433 ha), suggests that larger pathogen populations and more frequent use of fungicides from the QoI group could be responsible for the higher A143 allele frequency observed in the samples from Conara.^39^ Recently, the G143A mutation has been found at high frequencies in the Yorke Peninsula region of South Australia following new QoI control failure episodes in 2019. The deployment of the in-field G143A ASqPCR method during a field day allowed to correlate higher frequencies of the A143 allele in fields displaying high disease levels after QoI treatment. The results of the analysis were provided on site and the fact that many participants decided to review their chemical control strategies highlights the power of this approach not only as an in-season tool for the management of fungicide resistance, but also as part of more broader extension efforts.

We believe that the ASqPCR conducted in the field is a reliable tool that can be used for rapid and on-site quantification of *Bgt* mutants carrying the G143A Cytb mutation. An additional advantage of this method is that it can be adapted to detect changes in other genes encoding different fungicide targets. To that effect, a similar assay for the detection of pyrimethanil resistance in the grape pathogen *Botrytis cinerea* has already been trialled in the field (F. Lopez-Ruiz, personal communication).

The technology we describe here substantially increases the speed with which quantitative pathogen diagnosis and fungicide resistance evolution can be measured and delivered to growers. These techniques promise to markedly increase the flexibility of growers to modify fungicide timing, dose and product choices within a few hours of sampling.

The methodology could be enacted using portable instruments housed in the cab of a vehicle. However, we recognise that the true value of rapid diagnosis will be realised only when it is coupled with reliable advice and product choices. As such, the technology is perhaps best suited to implementation via testing stations located within cropping zones where information from the tests can be coupled with local pathology experience and weather forecasts. This would mean that the treatments used would be tailored to the disease and resistance problems within the crop, which in turn would eliminate the use of unnecessary treatments and reduce the selection pressure that occurs when using unsuitable treatments.

## Acknowledgements

This study was supported by the Centre for Crop and Disease Management, a co-investment of Curtin University and the Australian grains industry through the Grains Research and Development Corporation (research grant CUR00023). The authors wish to thank Rose Kristoffersen and Wesley Mair for the technical assistance during the Tasmanian and South Australian field trips, respectively. They also wish to thank Dr Ayalsew Zerihun for providing assistance with the Bland-Altman analysis, Foundation for Arable Research (FAR) for sample contribution and, Sam Trengove for coordinating sample submission at the field day in South Australia.

## Notes

### Competing Interest Statement

The authors have declared no competing interest.

